# *Magnaporthe oryzae SMO1* encodes a Ras GTPase-activating protein required for spore morphology, appressorium function and rice blast disease

**DOI:** 10.1101/388298

**Authors:** Michael J. Kershaw, Magdalena Basiewicz, Darren M. Soanes, Xia Yan, Lauren S. Ryder, Michael Csukai, Barbara Valent, Nicholas J. Talbot

## Abstract

The pathogenic life cycle of the rice blast fungus *Magnaporthe oryzae* involves a series of morphogenetic changes, essential for its ability to cause disease. The *smo* mutation was identified more than twenty-five years ago and affects the shape and development of diverse cell types in *M. oryzae,* including conidia, appressoria and asci. All attempts to clone the *SMO1* gene by map-based cloning and/or complementation, have failed over many years. Here, we report the identification of *SMO1* by a combination of bulk segregant analysis and comparative genome analysis. *SMO1* encodes a GTPase-activating protein (GAP), which regulates Ras signalling during infection-related development. Targeted deletion of *SMO1* results in abnormal, non-adherent conidia, impaired in their production of spore tip mucilage. Smo1 mutants also develop smaller appressoria, with a severely reduced capacity to infect rice plants. *SMO1* is necessary for organisation of microtubules and for septin-dependent remodelling of the F-actin cytoskeleton at the appressorium pore. Smo1 physically interacts with components of the Ras2 signaling complex, and a range of other signalling and cytoskeletal components, including the four core septins. *SMO1* is therefore necessary for regulation of RAS activation required for conidial morphogenesis and septin-mediated plant infection.

## Introduction

*Magnaporthe oryzae* (synonym of *Pyricularia oryzae)* is an ascomycete fungus responsible for rice blast disease (Zhang *et al.* 2016), a devastating plant disease resulting in severe losses to the global rice harvest each year. The need for increased rice production to feed the rapidly expanding human population, together with the increasing energy costs of both fungicides and fertilizers, means there is an urgent need to develop durable rice blast control strategies to be deployed as part of an environmentally sustainable plan for increasing global rice production (Wilson and Talbot, 2009; Yan and Talbot, 2016).

The rice blast fungus initiates plant infection when a three-celled spore, or conidium, lands and germinates on a leaf surface. Conidia are able to adhere to the hydrophobic leaf surface by means of spore tip mucilage, which is released from a compartment at the tip of the apical cell of the spore. Apical conidial attachment, together with the pyriform-shape of the spore, are hydrodynamically favourable for resisting water flow and maintaining attachment to the leaf as the spore germinates (Hamer *et al.* 1988). Typically, a single polarised germ tube emerges from the spore and after 4 to 6h, the tip of the germ tube swells and differentiates into a specialised infection cell called an appressorium (Ryder and Talbot, 2015; Talbot, 2003). In the appressorium, a discrete melanin cell wall layer is essential for generation of high internal turgor by facilitating accumulation of glycerol to very high concentrations (deJong *et al.* 1997). Penetration of the host cuticle results from the application of turgor as mechanical force, leading to protrusion of a rigid penetration peg to rupture the leaf cuticle. Re-polarisation of the appressorium requires septin mediated F-actin reorganisation at the base of the appressorium, where it contacts the leaf (Dagdas *et al.* 2012). The fungus invades host cells, colonizing tissue rapidly, which leads to formation of disease lesions from which the fungus produces large numbers of spores allowing rapid spread of the disease to neighbouring plants (Ou, 1985).

The *M. oryzae SMO1* locus was first defined from multiple mutants identified spontaneously, or through genetic screens that took place more than 25 years ago (Hamer *et al.* 1989b). One screen aimed to identify factors contributing to appressorium development and another involved isolation of mutants that were unable to adhere to hydrophobic surfaces, such as Teflon (poly-tetrafluoro-ethylene). All mutants formed aberrantly shaped spores, with no visible axis of symmetry. Wild-type conidia in *M. oryzae,* by contrast, are bilaterally symmetrical and pyriform (teardrop) shaped. These spore morphology mutants were named Smo and tetrad analysis showed that the phenotype was due to a single gene mutation that defined a new locus, *SMO1,* involved in cell shape determination. Smo1 mutants also developed misshapen asci and affected appressorium morphogenesis (Hamer et al. 1989b). The original *smo1* mutants were identified in a weeping lovegrass *(Eragrostis* curvula)-infecting *M. oryzae* strain 4091-5-8 (Hamer *et al.* 1989b), but *smo1* mutants were later isolated and characterised in a rice pathogen of *M. oryzae* and showed a virulence defect when inoculated on susceptible rice cultivars (Hamer and Givan, 1990). The *SMO1* locus was mapped based on the segregation of a dispersed repeated DNA sequence, called MGR586 (Hamer *et al.* 1989a), and shown to be located between two closely linked MGR sequences (Romao and Hamer, 1992). An exhaustive series of map-based cloning experiments and complementation analysis, however, failed to clone *SMO1,* so that its identity has remained unknown for the last 25 years.

Here, we report the identification of *SMO1* using comparative genome analysis and bulked segregant analysis. Complementation of the original Smo1 mutants, followed by targeted gene deletion confirmed the identity of *SMO1,* which encodes a GTPase-activating protein, most similar to *GapA* in *Aspergillus nidulans.* We show that *SMO1* is necessary for determination of conidial shape and the ability of spores to attach to hydrophobic substrates. Importantly, *SMO1* is also necessary for septin-mediated F-actin remodelling at the appressorium pore and therefore plays a critical role in plant infection by the rice blast fungus.

## Materials and Methods

### Fungal strains, growth conditions, and DNA analysis

*M. oryzae* strains used in this study were the rice pathogens Guy11 (Leung *et al.* 1988) and a *Δku-70* mutant impaired in non-homologous DNA-end joining (Kershaw and Talbot, 2009), the weeping lovegrass pathogen 4091-5-8, and 12 *smo1* mutants (see Table S1), either spontaneous mutants, or generated by UV mutagenesis in the original study (Hamer *et al.* 1989b). Growth, maintenance of *M. oryzae,* media composition, nucleic acid extraction, and transformation were all as described previously (Talbot *et al.* 1993b). Gel electrophoresis, restriction enzyme digestion, gel blots, DNA manipulation and sequencing were performed using standard procedures (Sambrook *et al.* 1989).

### Sequencing and Single Nucleotide Polymorphism analysis

Genomic DNA was extracted from each *smo1* mutant strain and sequenced using a HiSeq 2500 (Illumina, Inc.), generating 100 base paired-end reads. Reads were filtered using the fastq-mcf program from the ea-utils package (http://code.google.com/p/ea-utils/). Filtered reads were mapped against the *M. oryzae* (strain 70-15) reference genome version 8 (Dean *et al.* 2005) http://www.broadinstitute.org/annotation/genome/magnaporthe_comparative/) using Burrows-Wheeler Aligner (BWA;(Li and Durbin, 2009)). Bespoke Perl scripts were used to calculate mean aligned coverage of reads against the reference genome, to discover SNPs (based on minimum read depth of 10 and minimum base identity of 95%) and to identify genes in which SNPs occurred. Table S1 shows details of raw DNA sequence information generated from each mutant strain. Integrative Genome Viewer (IGV; (Robinson et al. 2011) was used to manually inspect read alignments for evidence of mutations in each strain. Sequence data for each mutant was submitted to the European Nucleotide Archive database (http://www.ebi.ac.uk/ena/data/view/PRJEB27449), and accession numbers are listed in Table S1.

### Bulked segregant genome analysis

Bulked segregant analysis (BSA) (Michelmore *et al.* 1991) was performed on a segregating ascospore population. Genetic crosses were performed, as described previously (Valent *et al.* 1991). Briefly, the two strains were inoculated together on oatmeal *agar* and grown at 24°C for 7 days and then at 20°C until flask-shaped perithecia were visible at the mycelial junction. Perithecia were transferred to 4% distilled water agar, separated from all conidia, and broken open to reveal asci. Mature asci were removed with a glass needle, and ascospores dissected from them. Ascospores were transferred individually to a 48-well plate containing complete medium and incubated for 4 to 5 days (Talbot *et al.* 1996). At this time, monoconidial re-isolations were made from each well and grown on individual plates for storage. Progeny were screened by microscopy and genomic DNA extracted from progeny using the CTAB method, described previously (Talbot *et al.* 1993a). DNA samples were bulked into two samples; wild type progeny and mutant progeny, and sequenced using HiSeq. Sequenced reads were aligned against the *M. oryzae* reference strain (70-15) assembly and examined for occurrence of SNPs segregating with the *smo1* mutation.

### Generation of targeted deletion mutants and strains expressing GFP fusions

Targeted gene replacement was carried out using a split-marker strategy (Catlett *et al.* 2003). Vectors were constructed using a hygromycin B resistance selectable marker, *hph* (Sweigard *et al.* 1997). To amplify split *hph* templates, the primers used were M13F with HY and M13R with YG, as described previously (Kershaw and Talbot, 2009). Sequence data for the *SMO1* candidate gene, MGG_03846, was retrieved from *M. oryzae* genome database at the Broad Institute (Massachusetts Institute of Technology, Cambridge, MA) (www.broad.mit.edu/annotation/fungi/magnaporthe) and used to design specific primer pairs (5’-Smo50.1/3’-SmoM13f/ and 5’-Smo30.1/3’-SmoM13r) to amplify regions flanking the open reading frame of MGG_03846 (Table S2). *M. oryzae* strain Guy-11 was transformed with the deletion cassettes (2 μg of DNA of each flank) and transformants selected in the presence of hygromycin B (200 μg ml^−1^). Two independent deletion mutants were obtained, as assessed by Southern blot analysis. A translational C-terminal MGG_03846 GFP fusion construct was generated by in-fusion cloning based on *in vitro* homologous recombination (Clontech). The primers 5’-Smop and 3’-SmoGFP were used to amplify a 4.5 kb fragment which included 1.9 kb of the MGG_03846 promoter region and 2.6 kb of the MGG_03846 open reading frame minus the stop codon. A 1.4 kb GFP fragment with trpC terminator, was amplified using primers 5’-smoGFP and 3’-TrpC, as listed in Table S2. Amplicons were cloned into pCB1532 (Sweigard *et al.* 1997), linearized with *BamHI* and *HindIII,* which carries the *ILV1* cassette conferring resistance to sulfonylurea. Homologous recombination results in assembly of fragments (5.9 kb) in the correct orientation to generate a gene fusion construct of 11.2kb. The construct was transformed into the wild type strain Guy11. For complementation of a *Δsmo1* mutant, the *SMO1-GFP* fusion cassette was transformed into the *Δsmo1-3* deletion mutant. A full length 5.6 kb fragment of the *SMO1* gene from Guy11 was amplified with primers 5’-Smop and 3’-smo30.1 (Table S2) and cloned into pCB1532 (Sweigard, 1997) and this construct was transformed into the *smo1* mutant CP751 (Hamer *et al.* 1989b). Transformants were selected in the presence of sulfonylurea (50 μg ml^−1^). For localisation of fluorescent fusion proteins in the *Δsmo1* mutant, *SEP3-GFP* (Dagdas *et al.* 2012), *Gelsolin-GFP* (Ryder *et al.* 2013) *Lifeact-GFP* (Berepiki *et al.* 2010), *GFP-ATG8* (Kershaw and Talbot, 2009) *H1-RFP* (tdTomato) (Saunders *et al.* 2010) and *β-tubulin:sGFP* (Saunders *et al.* 2010) constructs were transformed into *Δsmo1* and transformants selected on either sulfonylurea (50 μg ml^−1^) or bialophos (50 μg ml^−1^).

### Appressorium development, penetration assays and rice infections

*M. oryzae* conidia were obtained by harvesting suspensions in water from the surface of 12-day-old plate cultures prepared on CM agar. Infection-related development was assessed by incubating conidia on hydrophobic glass coverslips and allowing appressoria to form, before visualisation by epifluorescence or laser confocal microscopy. To visualise spore tip mucilage, FITC conjugated fluorescein isothiocyanate-conjugated concanavalin (FITC-ConA) was added at 1μgml^−1^ to harvested conidia and incubated at 24°C for 20 min before examination. Rice leaf sheath *(Oryza sativa)* inoculations were performed, as described previously, using the susceptible rice cultivar CO-39 (Kankanala *et al.* 2007). Appressorium-mediated penetration of onion *(Allium cepa)* epidermal strips was assayed, as described previously (Balhadere *et al.* 1999) and assessed by recording the frequency of hyphal penetration from an appressorium. An incipient cytorrhysis assay was carried out by allowing appressoria to form in water on borosilicate cover slips for 24 h after which the water was replaced with a range of aqueous glycerol ranging from 0.25M to 2.5M and after 30 min, the frequency of cytorrhysis determined (de Jong *et al.* 1997). Plant infection assays were performed by spraying seedlings of rice cultivar CO-39 with a suspension of 10^5^ conidia per ml^−1^, as previously described (Talbot *et al.* 1993b). Occurrence of blast symptoms was recorded 5 days after inoculation and experiments performed three times.

### Protein-protein interaction studies

A yeast two-hybrid screen was performed to determine physical interactions of Smo1 and to investigate its function as a Ras-GAP, using the Matchmaker GAL4 Two-Hybrid system 3 (Clontech Laboratories, Inc.). *SMO1, RAS2,* and *RAS1* cDNA were cloned into the bait vector pGBKT7 using primer combinations, 5’-smoGB/ 3’-smoGB, 5’-ras2GB/3’ras2GB and 5’-ras1GB/ras1GB. *SMO1, RAS2,* and *GEF1* cDNA were cloned into the prey vector PGADT7 using primer combinations 5’-smoGA/ 3’-smoGA, 5’-ras2GA/3’ras2GA and 5’-gef1 GA/gef 1GA. Cloning was performed using in-fusion cloning (Clontech Laboratories, Inc.). Sequencing was performed to ensure constructs were in-frame (MWG operon). Yeast two-hybrid analysis was then carried out using the Matchmaker GAL4 Two-Hybrid system 3 (Clontech), according to the manufacturer’s instructions (Wilson *et al.* 2010). For *in vivo* co-immunoprecipitation studies, total protein was extracted from lyophilized *M. oryzae* mycelium of strains expressing Smo1:GFP and ToxA:GFP (control) after growth in liquid CM for 48 h. Protein extracts were co-immunoprecipitated using the GFP-Trap protocol, according to the manufacturer’s protocol (ChromoTek). Peptides were prepared for LC-MS/MS and separated by SDS/PAGE. Gels were cut into slices and LC-MS/MS analysis was performed at the University of Bristol Proteomics Facility.

### Light and Epifluorescence Microscopy

Epifluorescence microscopy was used to visualize localisation of fluorescent fusion proteins expressing eGFP or RFP using a IX81 motorized inverted microscope (Olympus) equipped with a UPlanSApo 100X/1.40 Oil objective (Olympus). Excitation of fluorescently-labelled proteins was carried out using a VS-LMS4 Laser-Merge-System with solid-state lasers (488 nm/50 mW). Laser intensity was controlled by a VS-AOTF100 System and coupled into the light path using a VS-20 Laser-Lens-System (Visitron System). Images were captured using a Charged-Coupled Device camera (Photometric CoolSNAP HQ2, Roper Scientific). All parts of the system were under the control of the software package MetaMorph (Molecular Devices, Downingtown, USA). Supplemental Movie 1 was performed using a Leica SP8 laser confocal microscope, with an argon laser line (488 nm) to excite GFP for imaging.

## Results

### Identification of the SMO1 locus

To identify the *SMO1* locus, we carried out bulked segregant analysis (Michelmore *et al.* 1991) using whole genome sequencing to identify single nucleotide polymorphisms. *M. oryzae* strain 4395-4-4 *(smo1 alb1 Mat1-2)* was crossed with a wild type strain TH3 *(Mat1-1)* and 50 ascospores collected. Ascospore progeny were phenotypically characterised based on spore shape, as *smo1* mutants have one, or two-celled spherical, or misshapen conidia, compared to the three-celled pyriform wild-type conidia. Progeny were therefore selected according to the Smo1 phenotype, DNA extracted from each individual and then bulked. Whole genome sequencing of bulked DNA samples identified a region of 2,061,034 bases on supercontig 8.6, which was defined by SNPs showing greater than 85% linkage to *SMO1* using bulk segregant analysis (BSA) (Figure 1A). This conformed to the region originally defined by MGR586-based genetic mapping spanning the *SMO1* locus (Romao and Hamer, 1992). Consistent with this, two single copy RFLP probes previously shown to be closely linked to the *SMO1* locus (Hamer and Givan, 1990; Romao and Hamer, 1992): JH4.28 and JH5.00, were sequenced and mapped to the *M. oryzae* genome. They were both found on supercontig 8.6, separated by a distance of 1,502,200 bases. The identified *SMO1* gene lies within this region, as identified by BSA (Figure 1A).

**Figure 1.**
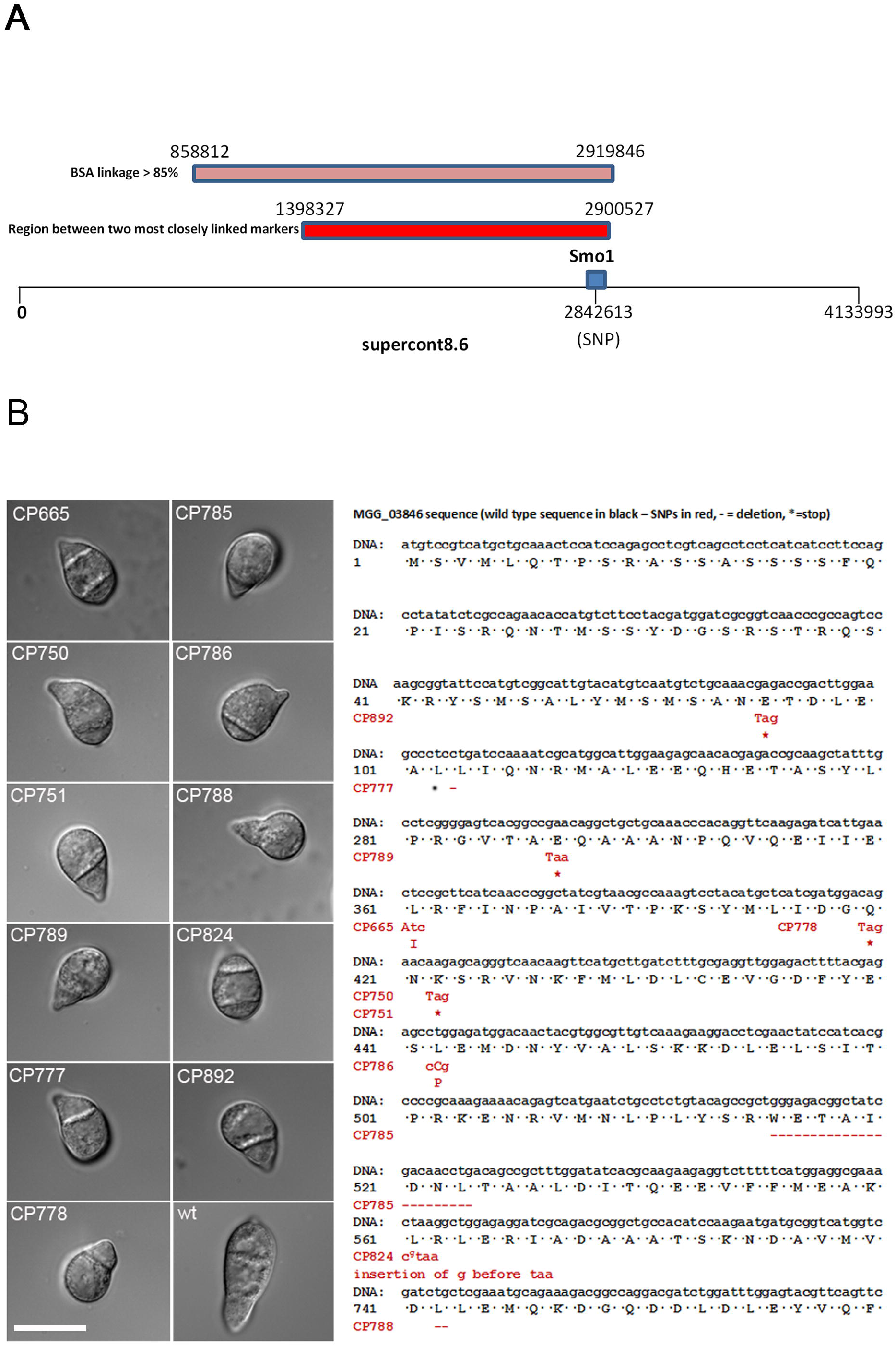
Identification of the *SMO1* locus in *Magnaporthe oryzae.* **A**. The identified *SMO1* gene lies on supercontig 8.6 (Chromosome 6: 2,838,047-2,844,822, ensembl database). Single copy RFLP probes JH4.28 and JH5.00, which have previously been shown to be closely linked to the *SMO1* locus (Hamer and Givan, 1990; Romao and Hamer, 1992), were sequenced and mapped to the *M. oryzae* genome, and also found on supercontig 8.6, separated by a distance of 1,502,200 bases. A region in a similar position on supercontig 8.6 was defined by SNPs that showed a greater than 85% linkage to *SMO1* using bulk segregant analysis (BSA). **B**. Micrographs showing spore morphology of *smo1* strains used in this study compared to wild type strain Guy11 (Bars = 10μm). Smo1^−^ mutants were originally obtained spontaneously or after UV mutagenesis (Hamer *et al.* 1989b). Nucleotide and amino acid sequences of MGG_03846 indicating position of SNPs and subsequent mutation in each of the *smo1* mutants. Wild type sequence in black, SNPs in red, * = stop codon.

We then analysed an allelic series of original *smo1* mutants, which were either selected by UB mutagenesis as non-adherent mutants, appressorium development mutants, or spontaneous Smo1 mutants (Hamer *et al.* 1989b), as described in Table S1, and confirmed their phenotype by spore morphology, as shown in Figure 1B. Genomic DNA was extracted from each strain and whole genome sequencing carried out. We also carried out genome sequence analysis of the parental 4091-5-8 strain, from which the Smo1 mutants were originally selected (Hamer *et al.* 1989b). All Smo1 mutants shared 202,020 SNPs which distinguished 4091-5-8 from the *M. oryzae* genome reference strain 70-15. Subtracting these shared SNPs from the derived SNP datasets for each strain, identified SNPs unique to each mutant. Six of the ten mutant *smo1* strains possessed a SNP in gene MGG_03846. Manual inspection of reads aligned to MGG_03846 identified mutations in five other mutant strains (Figure 1B). Only CP790 did not contain a mutation in MGG_03846, and our analysis suggests that this strain does not exhibit the Smo1 phenotype and therefore may have reverted since the original study (Hamer *et al.* 1989b). With the exception of strains CP750 and CP751, which had identical SNPs, the mutations were different in each strain, consistent with the allelic variability reported originally (Hamer *et al.* 1989b). Five of the strains had a SNP which introduces a stop codon into the open reading frame (nonsense mutation), four had either an insertion or deletion that produces a frameshift mutation and two possessed a base pair substitution that resulted in a change in an amino acid residue (Figure 1B). The mutation in CP786, for instance, changes a leucine to proline, which is likely to produce a distinct alteration in secondary structure, given that proline acts as a structural disruptor in the middle of regular secondary structure elements, such as alpha helices and beta sheets (Williamson, 1994).

### Cloning and characterisation of the SMO1 gene of M. oryzae

The *SMO1* candidate gene, MGG_03846, is 2643 bp in length with four introns of 81, 83, 78 and 65 bp, respectively, and encodes a putative 780 aa protein. Bioinformatic analysis predicted that MGG_03846 encodes a Ras GTPase activating protein (RasGAP), and the predicted gene product possesses two domains, a GTPase activator domain for Ras-like GTPase from amino acids 193-401 and a RAS GAP C-terminal motif from amino acids 580-699. There are four putative GapA encoding genes in *M. oryzae* and phylogenetic analysis (Figure S1) revealed potential *SMO1* orthologs of *GapA* from *Aspergillus nidulans* (Harispe *et al.* 2008) and *Gap1* from *Schizosaccharomyces pombe* (Imai *et al.* 1991). Phylogenetic analysis of other putative GAPs in *M. oryzae* suggests that MGG_03700 is a homolog of the *Saccharomyces cerevisiae* IQG1, which controls actin-ring formation and cytokinesis (Epp and Chant, 1997). MGG_08105 is a homolog of the S. *cerevisiae BUD2,* which plays a role in spindle position checkpoint and bud site selection (Park *et al.* 1993), whilst MGG_11425 is a homolog of the S. *cerevisiae* RasGAPs *IRA1* and *IRA2,* which are negative regulators of Ras-cAMP signaling pathway required for reducing cAMP levels under nutrient limiting conditions (Tanaka *et al.* 1989; Tanaka *et al.* 1990) (Figure S1).

To determine whether the candidate RAS-GAP-encoding gene is *SMO1* we cloned a full length copy of MGG_03846 under control of its native promoter and transformed this into *smo1* mutant, CP750 (Hamer *et al.* 1989b). Re-introduction of *SMO1* fully complemented the *smo1* spore shape phenotype, and spores in the complemented strain germinated to produce short germ tubes and normal appressoria, which were able to cause rice blast disease (Figure S2).

### Targeted deletion of SMO1 leads to cell shape defects

We next carried out targeted deletion to generate a *Δsmo1* mutant and two independent deletion strains were selected and confirmed by Southern blot analysis for further analysis. The morphology of *Δsmo1* mutants was similar to the original *smo1* mutants, with mycelial colonies more compact, white and fluffy, compared to those of the isogenic wild type strain Guy-11. Conidia were more rounded and predominantly unicellular, or two-celled (Figure 2A). By contrast, wild type spores are typically 22-25μm in length, and 8.75 and 5.5μm in diameter, whereas *Δsmo1* conidia are typically 12.5-15 μm in length, and 8.5-10 μm in diameter. Conidia of *Δsmo1* mutants germinate normally, but produce longer germ tubes than the wild type, typically 87μm in *Δsmo1* compared to 22μm in Guy11 (Figure 3A). Appressorium development was delayed and appressoria were typically misshapen and slightly smaller, 8.64 ± 0.56μm in diameter in *Δsmo1,* compared to 9.55 ± 0.56μm in Guy11 (p < 0.01), as shown in Figure 3.

**Figure 2.**
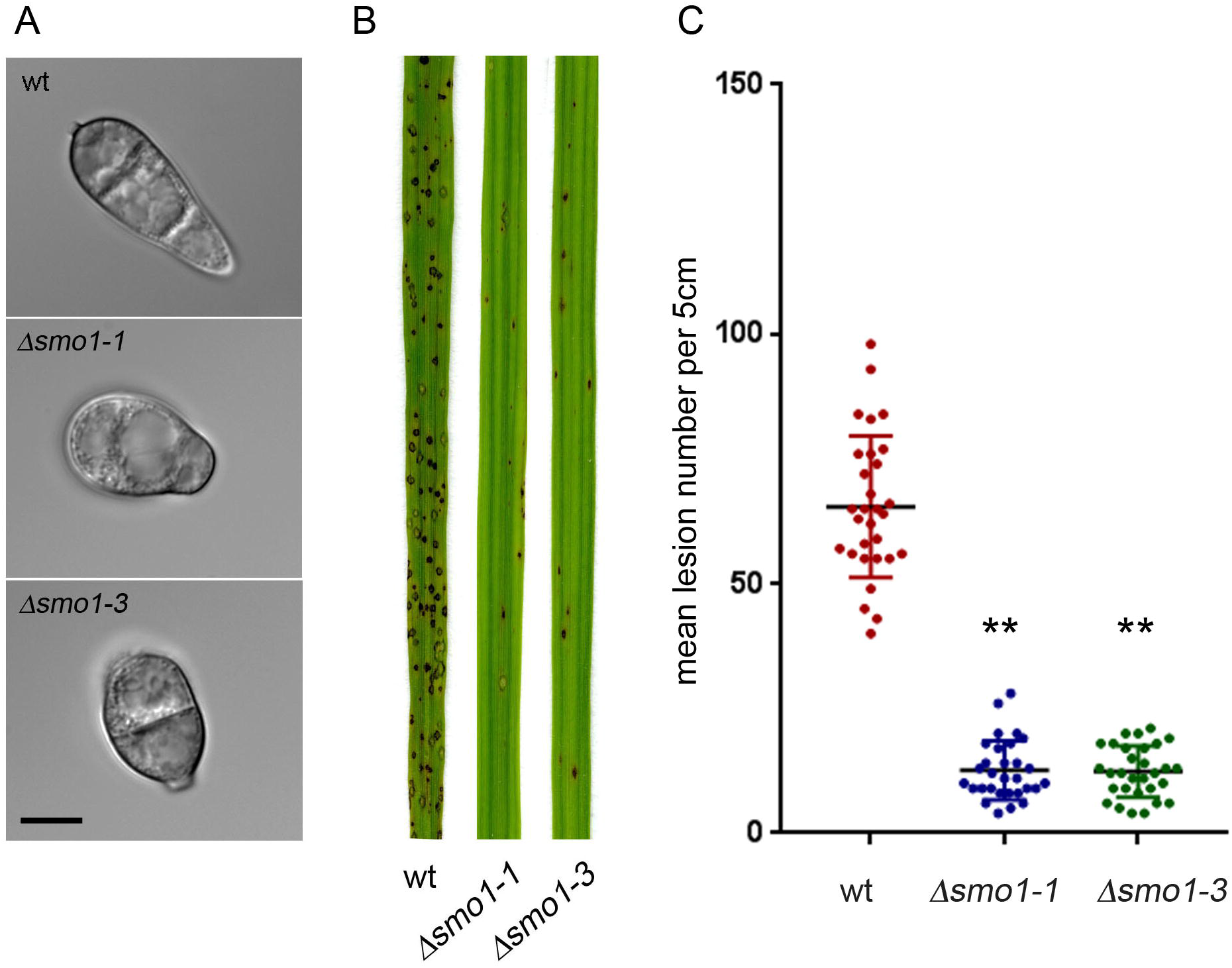
Deletion of MGG_03846 resulted in *smo1* spore phenotype and mutants were reduced in plant infection. **A**. Micrographs showing spores of two MGG_03846 deletion strains *(Δsmo1)* as compared to wild type strain Guy11 (Bar = 5μm). **B**. Photographs of leaves from infected plants. Seedlings of rice cultivar CO-39 were inoculated with conidial suspensions (5 × 10^4^ ml^−1^). Seedlings were incubated for 5 days to allow symptom development. **C**. Box plot of mean lesion density per 5 cm leaf after infection with MGG_03846 deletion mutants compared to wild type. Error bar equals standard error of the mean. * *P* < 0.0001 (unpaired Students’s *t* test (n = 3 experiments of 40 leaves).

**Figure 3.**
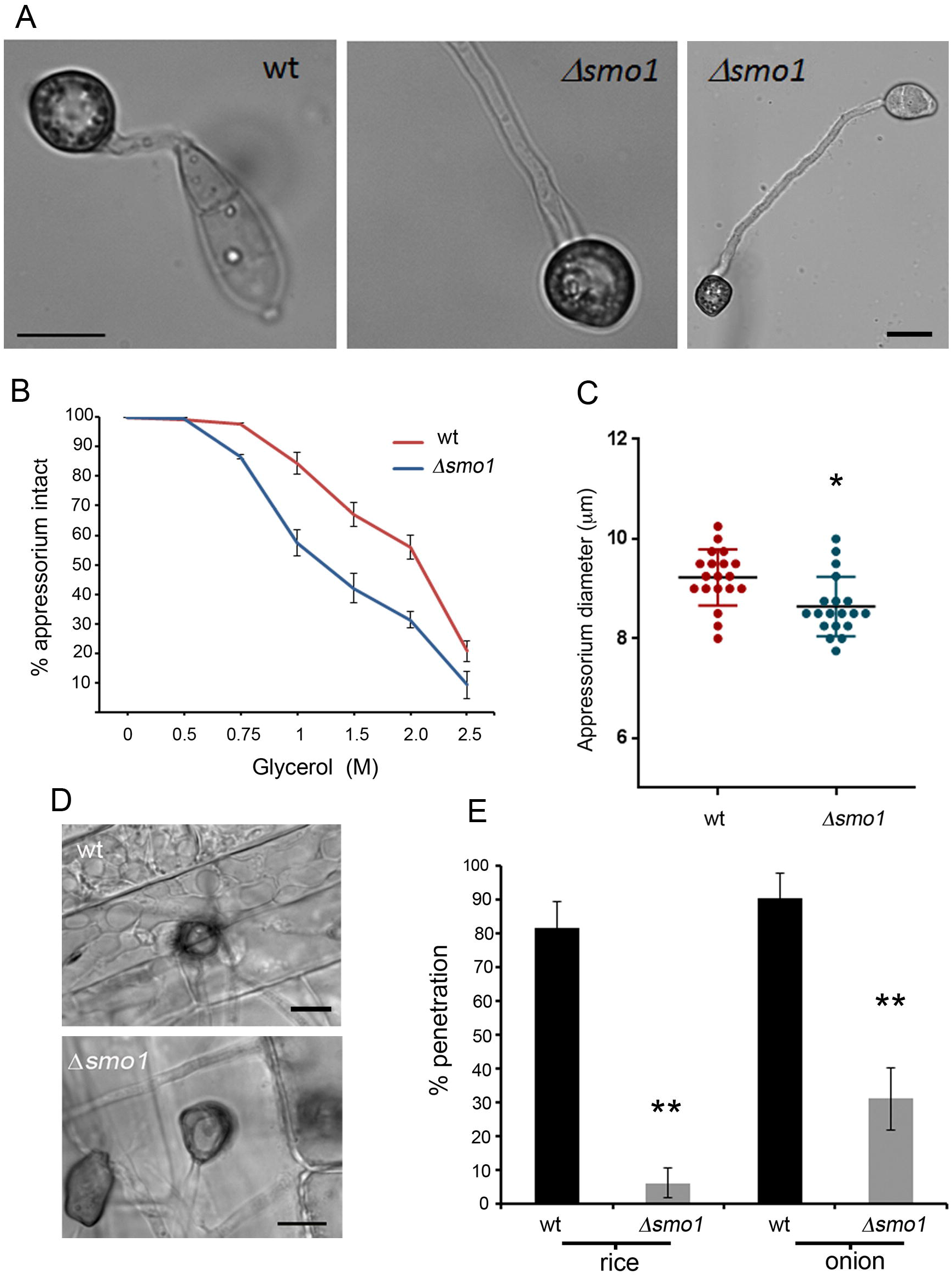
The *Δsmo1* mutant is able to elaborate an appressorium, which is impaired in function. **A**. Micrograph showing *Δsmo1* mutant with extended germ tube compared to wild type strain (Bar = 5μm). **B**. Incipient cytorrhysis assays measuring intracellular glycerol were carried out by allowing appressoria to form in water on borosilicate cover slips for 24 h, after which the water was replaced with a range of aqueous glycerol solutions ranging from 0.25 M to 2.5 M. The rate of cell collapse was determined after 30 min. A concentration of 1.25 M glycerol caused the collapse of 50% of appressoria in the *Δsmo1* mutant whereas 2.25 M glycerol was required for 50% collapse in the wild type. *P* < 0.0001 unpaired Student’s *t* test, n = 100). **C**. Box plot showing reduced appressorium diameter of *Δsmo1* mutant compared to wild type, Guy11. \**P* < 0.01 unpaired Student’s *t* test, n = 50. **D**. Micrograph comparing penetration of rice leaf sheath by Guy11 and the *Δsmo1* mutant. Inoculations were performed as described previously using the susceptible rice line CO-39. **E**. Bar chart showing percentage penetration of Guy11 and *Δsmo1* mutant on leaf sheath and onion epidermis assessed by recording the frequency of hyphal penetration from an appressorium **P< 0.001 unpaired Student’s *t* test, n = 24.

As a consequence of the abnormal spore morphology and delay in appressorium formation in *Δsmo1* mutants, we decided to investigate the pattern of nuclear division during appressorium development. In *M. oryzae,* a single round of mitosis occurs prior to appressorium development, followed by conidial cell death and degradation of nuclei in each conidial cell (Veneault-Fourrey *et al.* 2006; Saunders *et al.* 2010). We introduced an H1-GFP fusion into the *Δsmo1* mutant to visualise nuclear dynamics by live-cell imaging (Figure S3A). Nuclear division in *Δsmo1* takes place within 4-6 hours post inoculation (hpi), as observed in Guy11 and, in addition, one daughter nucleus migrates to the developing appressorium and the other nucleus returned to the conidium, in the same way as the wild type (Figure S3A). Nuclear material was, however, often observed in the longer germ tube, as well as the conidium. We used Calcofluor-white to examine septation events in the germ tube and observed that one septum normally forms in the germ tube, often near to the conidium (Figure S3B). After 16 h, nuclei in *Δsmo1* conidia start to degrade and by 24 h the spore had collapsed, as was observed in Guy11 (Figure S3A). We conclude that *Δsmo1* mutants show defects in spore shape and organisation and exhibit extended germ tube growth associated with a delay in appressorium development.

### *The* SMO1 *gene encodes a virulence factor in* M. oryzae

To determine the role of *SMO1* in fungal pathogenicity, we inoculated the susceptible rice cultivar CO-39 with spore suspensions of two *Δsmo1* mutant strains and Guy11.

The *Δsmo1* mutants generated significantly reduced numbers of disease lesions, 13.44 ± 1.34 per 5 cm leaf, compared to 65 ± 6.94 lesions per 5 cm leaf in Guy11 (P<0.0001) (Figure 2B & C). To determine whether the reduced ability of *Δsmo1* mutants to cause disease lesions was due to reduced appressorium turgor, we incubated appressoria of *Δsmo1* mutants in a series of glycerol solutions of increasing molarity and measured the frequency of incipient cytorrhysis (cell collapse) (Howard *et al.* 1991). A concentration of 1.25 M glycerol caused the collapse of 50% of appressoria in the *Δsmo1* mutant whereas in the wild type 2.25 M glycerol was required for 50% (*P* < 0.0001) (Figure 3C). This reduction in turgor is consistent with an appressorium penetration defect. We therefore applied the *Δsmo1* mutant to excised rice leaf sheath to determine the frequency of appressorium-mediated penetration. We observed that Guy11 had a frequency of successful penetration events of 81.66% ± 7.59 penetration, compared to 6.3 ± 4.37 in *Δsmo1* (P<0.001), as shown in Figure 3D & E. We also tested whether *Δsmo1* mutants were able to penetrate onion epidermis and found slightly increased penetration compared to that of *Δsmo1* mutants on rice leaf sheaths, but still significantly reduced compared to the wild type (90.33. ± 7.59 (wt) compared to 31 ± 9.23 in *Δsmo1; P* < 0.001), as shown in Figure 3E. We conclude that *Δsmo1* mutants are reduced in their ability to cause rice blast disease because of impairment in appressorium function, including a reduction in turgor and frequency of penetration peg development.

### Smo1 localises to the appressorium pore during plant infection

To determine the subcellular localisation of the Smo1 protein and its temporal dynamics during infection-related development, we generated a *SMO1-GFP* fusion which was introduced into Guy11 and a *Δsmo1* mutant. Expression of *SMO1-GFP* in the *Δsmo1* mutant strain was sufficient to fully complement the phenotypes associated with the *Δsmo1* mutant, restoring wild type spore and appressorium morphologies and the ability to infect rice and cause disease (Figure 4 & Figure S2). Analysis of the cellular localisation showed that Smo1 localises to the tip of germ tubes during germination. As the appressorium forms, Smo1 localised initially as small puncta throughout the appressorium (Figure 4A). However, after 24 h when maximal turgor is established in the appressorium, Smo1 localisation became more condensed and by using 3-D reconstruction of a mature appressorium, Smo1-GFP was observed to localise predominantly to the base of the appressorium around the appressorium pore (Figure 4B, Supplemental Movie 1). Smo1 distribution is therefore associated with regions of polarised growth, such as the germ tube tip. In the appressorium, Smo1 localises to the point at which anisotropic growth is reestablished for penetration peg development and plant infection.

**Figure 4.**
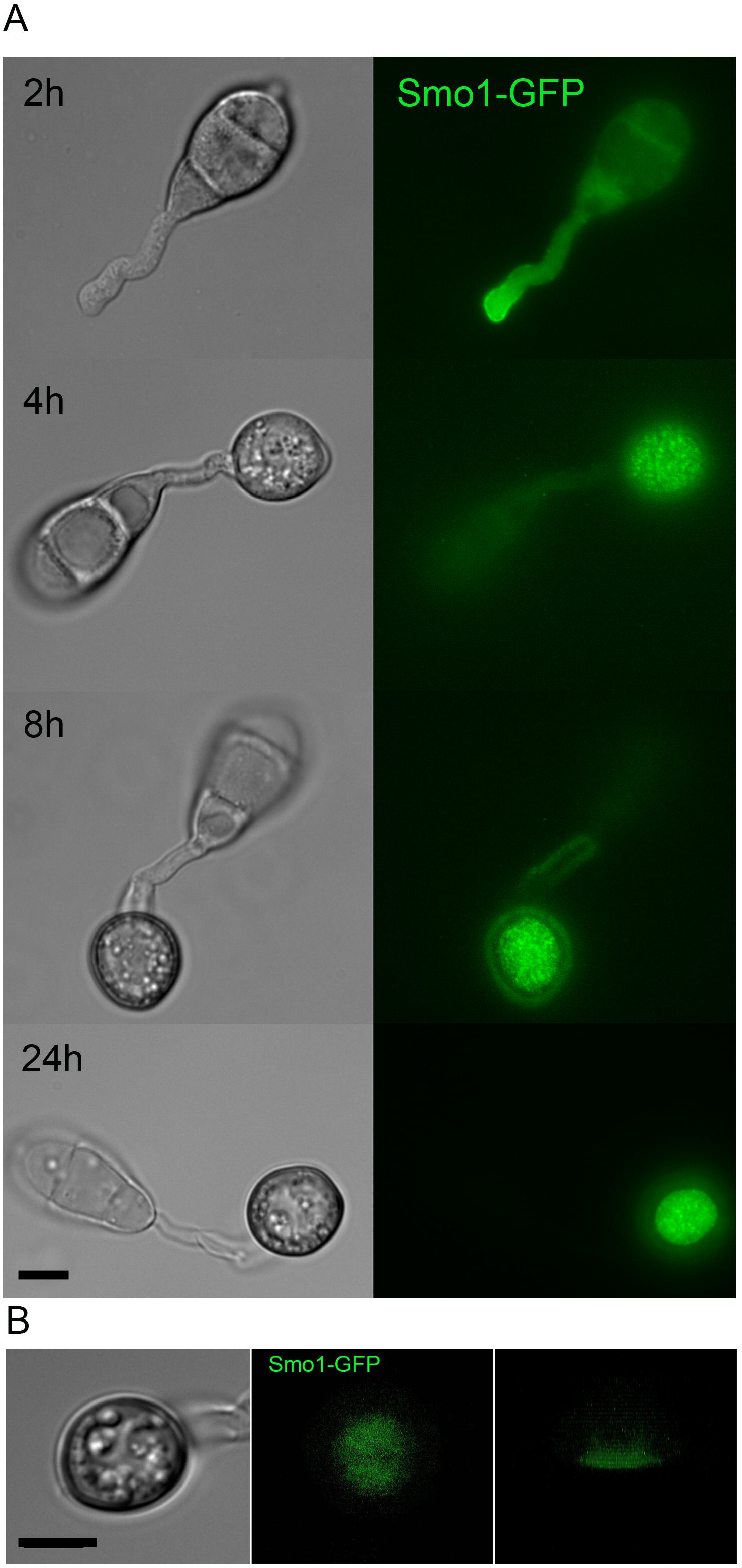
Live cell imaging of *M. oryzae* wild type strain expressing Smo1-GFP **A**. Cellular localisation of Smo1-GFP at the tip of the germ tube and as punctate structures in both the developing appressoria **B**. Distribution of Smo1-GFP in mature appressorium visualised by laser confocal miscoscopy. The right hand panel shows confocal image of transverse view of appressorium with Smo1-GFP present predominantly at the appressorium-substrate interface (see also Supplemental Movie 1). Spores were harvested and inoculated onto hydrophobic cover slips and visualised by epifluorescence or laser confocal microscopy. Bar = 10 μm.

### Δsmo1 *mutants are defective in spore tip mucilage generation and surface attachment*

The tight adhesion of conidia to the rice leaf surface is critical for rice blast disease and involves release of spore tip mucilage (STM) from the tip of the conidium (Hamer *et al.* 1988). We used FITC-labelled concanavalin-A (ConA-FITC) to compare STM released from spores of Guy-11 and the *Δsmo1* mutant. We evaluated levels of STM secretion at 30 min and then 2 h post-inoculation. This revealed a clear reduction in STM secretion in *Δsmo1* mutants compared to Guy-11. During the early stages of conidial attachment and germination, STM in the *Δsmo1* mutant was noticeably reduced compared to Guy-11 (Figure 5A & 5B). We also observed Guy-11 and *Δsmo1* mutants after 24 h and observed a similar reduction in a ConA-positive mucilage layer around the mature appressorium in a *Δsmo1* mutant (Figure 5C). To examine conidial adhesion, we then counted the number of conidia that could be removed from the surface of hydrophobic coverslips by washing 30 min after inoculation. In Guy11, 67 ± 6.9% of conidia remained attached to PTFE Teflon surfaces after washing, whereas in *Δsmo1* mutants 48.4 ± 9.97% (P < 0.01) remained attached (Figure 5D). We conclude that STM secretion is impaired in *Δsmo1* mutants which show reduced adhesion to hydrophobic surfaces.

**Figure 5.**
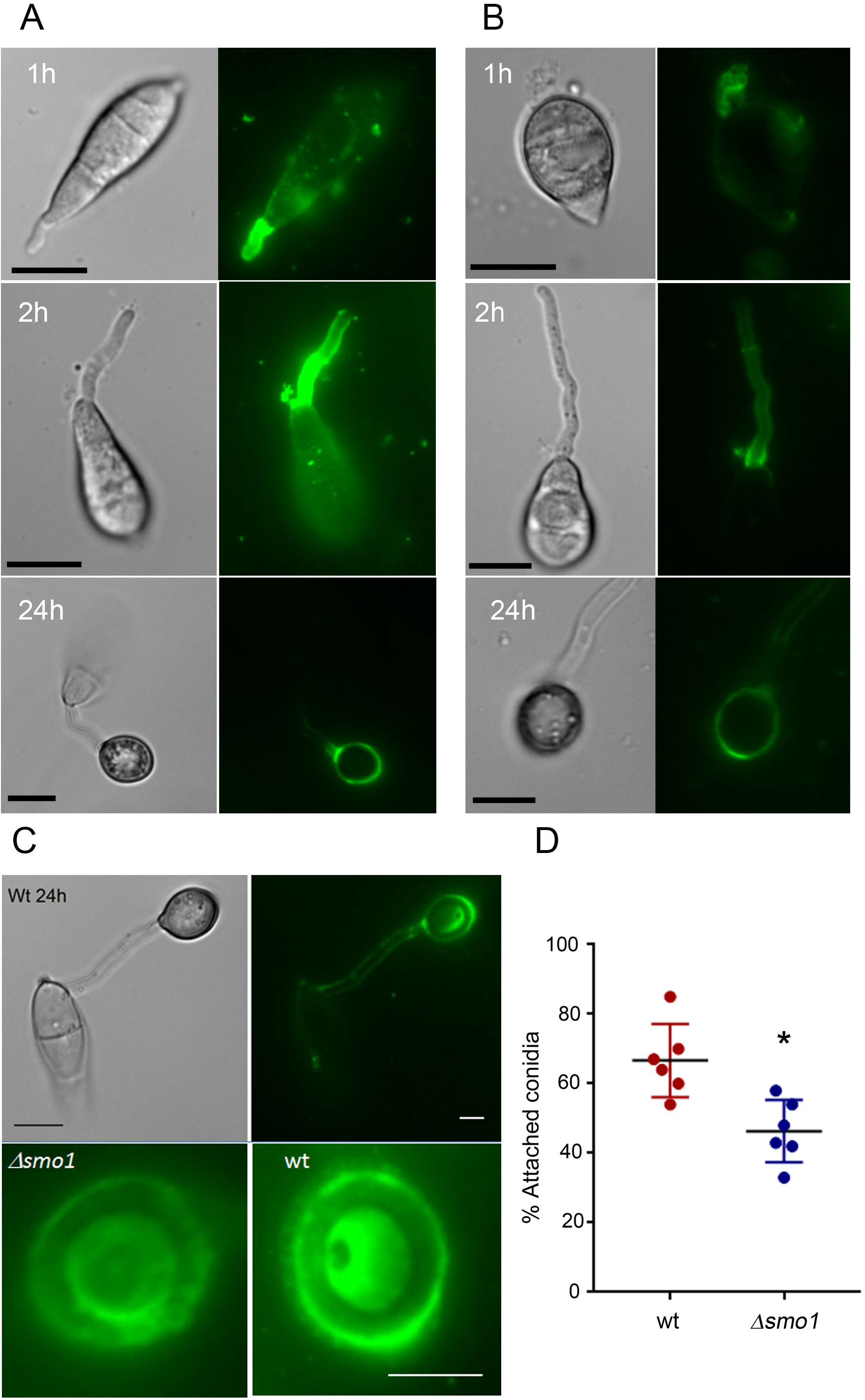
Live cell imaging of *M. oryzae* wildtype and *Δsmo1* mutant showing release and levels of spore tip mucilage (STM) by addition of ConA FITC to germinating *M. oryzae conidia* of (A) Guy11 and (B) *Δsmo1* mutant. Spores were harvested and inoculated onto hydrophobic cover slips before addition of ConA FITC (100 ug/ml). Bar = 10μm. (C) ConA FITC staining of mucilage in mature appressoria of Guy11 and *Δsmo1* mutant after 24 h. Bar = 5μm. (D) Bar chart showing percentage of germinated conidia of Guy11 and *Δsmo1* remaining attached to cover slips after washing. Conidia were harvested and counted (1 × 10^5^ ml^−1^) and allowed to attach to hydrophobic cover slips 30 minutes before washing with water. *P< 0.01 unpaired Student’s *t* test, n = 100.

### Δsmo1 *mutants are impaired in septin-mediated F-actin reorganisation at the appressorium pore*

The conidial shape phenotype of *Δsmo1* mutants suggested an effect on the distribution and organisation of cytoskeletal components. We therefore visualised the distribution of microtubules based on expression of the β-tubulin Tub2-GFP fusion protein. Conidia of the wild type Guy11 showed a network of long microtubules defining each of the three-cells within the spore (Figure 6A and Supplemental Movie 2). By contrast microtubules observed in spores of a *Δsmo1* mutant showed an abnormal distribution, consistent with the spherical shape of spores (Figure 6A and Supplemental Movie 3). A key requirement for appressorium function in *M. oryzae* is the recruitment and organisation of a septin-dependent toroidal F-actin network at the appressorium pore. Septins provide cortical rigidity to the infection cell at the point of penetration and act as a diffusion barrier for organisation of polarity determinants required for penetration hypha development (Dagdas *et al.* 2012). We decided to investigate organisation of the septin Sep5-GFP (Dagdas *et al.* 2012) in the *Δsmo1* mutant. In Guy11, a septin ring was visible surrounding the appressorium pore, but this was mis-localised in the *Δsmo1* mutant (Figure 6B). We therefore observed the F-actin cytoskeleton, by expressing actin-binding protein gene fusions LifeAct-RFP and Gelsolin-GFP (Berepiki *et al.* 2010; Ryder *et al.* 2013) in the *Δsmo1* mutants and observing appressoria at 24 hpi. In Guy-11, Lifeact-GFP and Gelsolin-GFP fluorescence revealed the toroidal F-actin network at the appressorium pore which marks the point at which the penetration peg emerges (Figure 6B). By contrast, the *Δsmo1* mutant showed dispersed and non-specific localisation of LifeAct-RFP and Gelsolin-GFP (Figure 6B). We conclude that the septin-mediated F-actin dynamics necessary for host penetration are regulated by signalling pathways acting downstream of Smo1.

**Figure 6.**
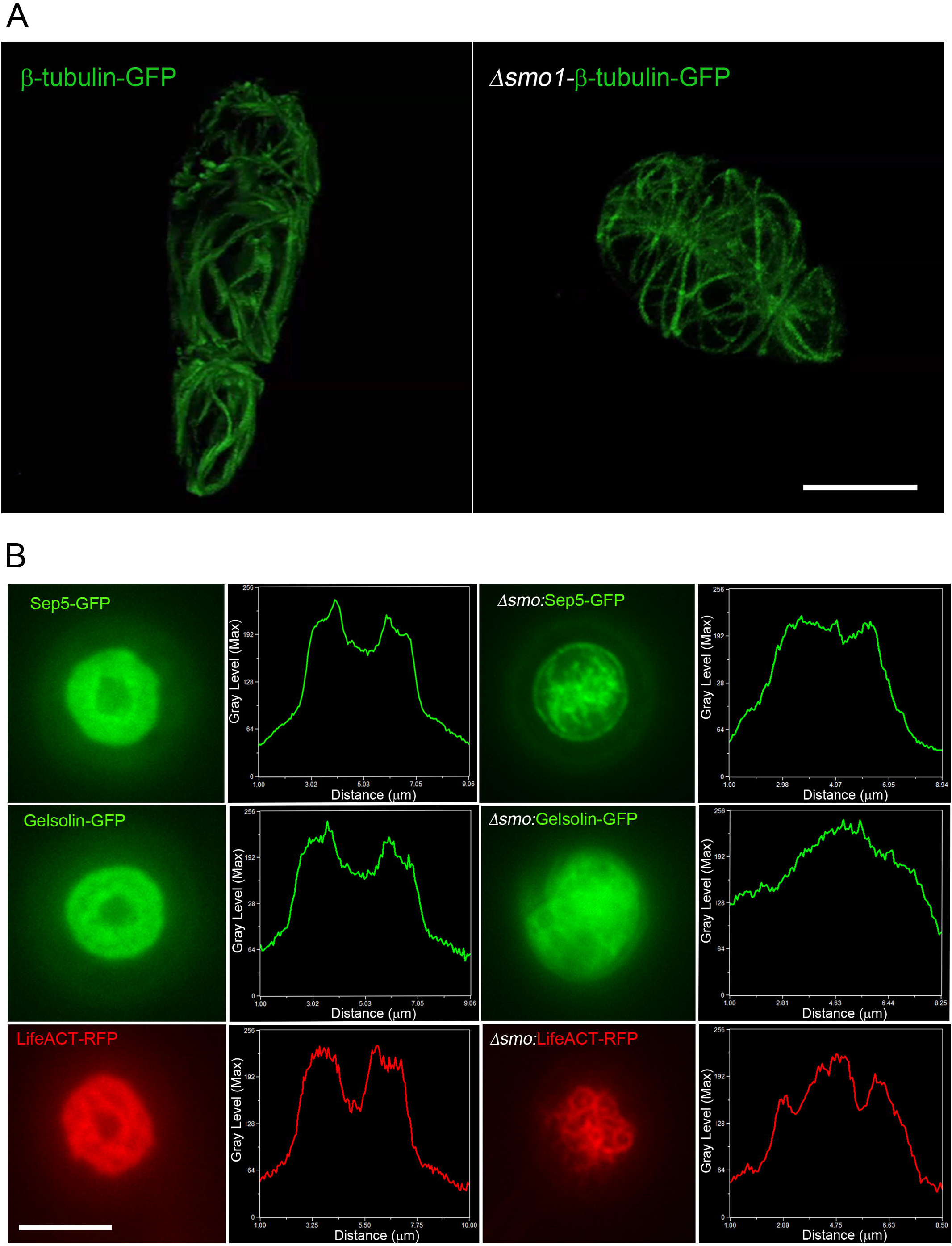
*Δsmo1* mutants are unable to undergo septin-mediated F-actin remodelling in the appressorium. (A) Expression of β-tubulin-GFP in conidia of *M. oryzae* Guy11 (left hand panel) and *Δsmo1* mutant (right hand panel). Microtubules showed aberrant distribution and organisation, consistent with spore-shape defect in *Δsmo1* mutant (B) Live cell imaging of septin-dependent F-actin network in appressoria of *M. oryzae* Guy11 and *Δsmo1* mutant at 24 hpi, visualised by laser-assisted epifluorescence microscopy. Localisation of Sep5-GFP, Gelsolin-GFP, and LifeAct-RFP at the appressorium pore with corresponding line-scan graphs to show distribution of fluorescence signal in a transverse section. Organisation of appressorium pore components requires Smo1. Bar = 5 μm.

### Protein-protein interaction studies to identify Smo1-interacting partners

The identification of Smo1 as a putative Ras-Gap protein prompted us to identify its potential interacting partners. Two independent lines of investigation were followed. First of all, yeast two hybrid analysis was carried out between Smo1 and confirmed Ras signalling components from *M. oryzae.* Initially, control experiments were performed in which the pGAD–Smo1 (prey–Smo1), pGAD–Ras2 (prey–Ras2), pGAD–Gef1(prey–Smo1), and also pGBK–Ras(bait-Ras2), pGBK–Smo 1 (bait— Smo1) and pGBK–Ras1 (bait—Ras1) were independently transformed into the yeast two hybrid Gold strain before plating onto SD/-Leu and SD/-Trp media, respectively. The lack of growth on these media demonstrates that none of the vectors are capable of auto-activating reporter genes. Simultaneous co-transformation of the pGBK-Ras2 (bait–Ras2) and pGAD–Smo1 (prey—Smo1) vectors into the Y2H Gold strain resulted in activation of all four reporter genes and growth on high stringency medium (-His/-Ade/-Leu/-Trp/+X-α-Gal) (Figure 7A). Co-transformation also activated *MEL1* expression in which the enzyme α–galactosidase is secreted into the medium, resulting in hydrolysis of X-α-Gal in the medium and turning the yeast colony blue. Growth on such high stringency media supports the hypothesis that Smo1 and Ras2 can physically interact. Putative interactions were also observed between Smo1 and Gef1, and between Ras2 and Gef1. Weaker interactions were also observed between Ras1 and both Gef1 and Smo1. When considered together, the interactions are consistent with Smo1 acting as a GTPase-activating protein on Ras2. We cannot preclude that Smo1 also plays a regulatory role in Ras1 signalling, but it shows much higher affinity to Ras2.

**Figure 7.**
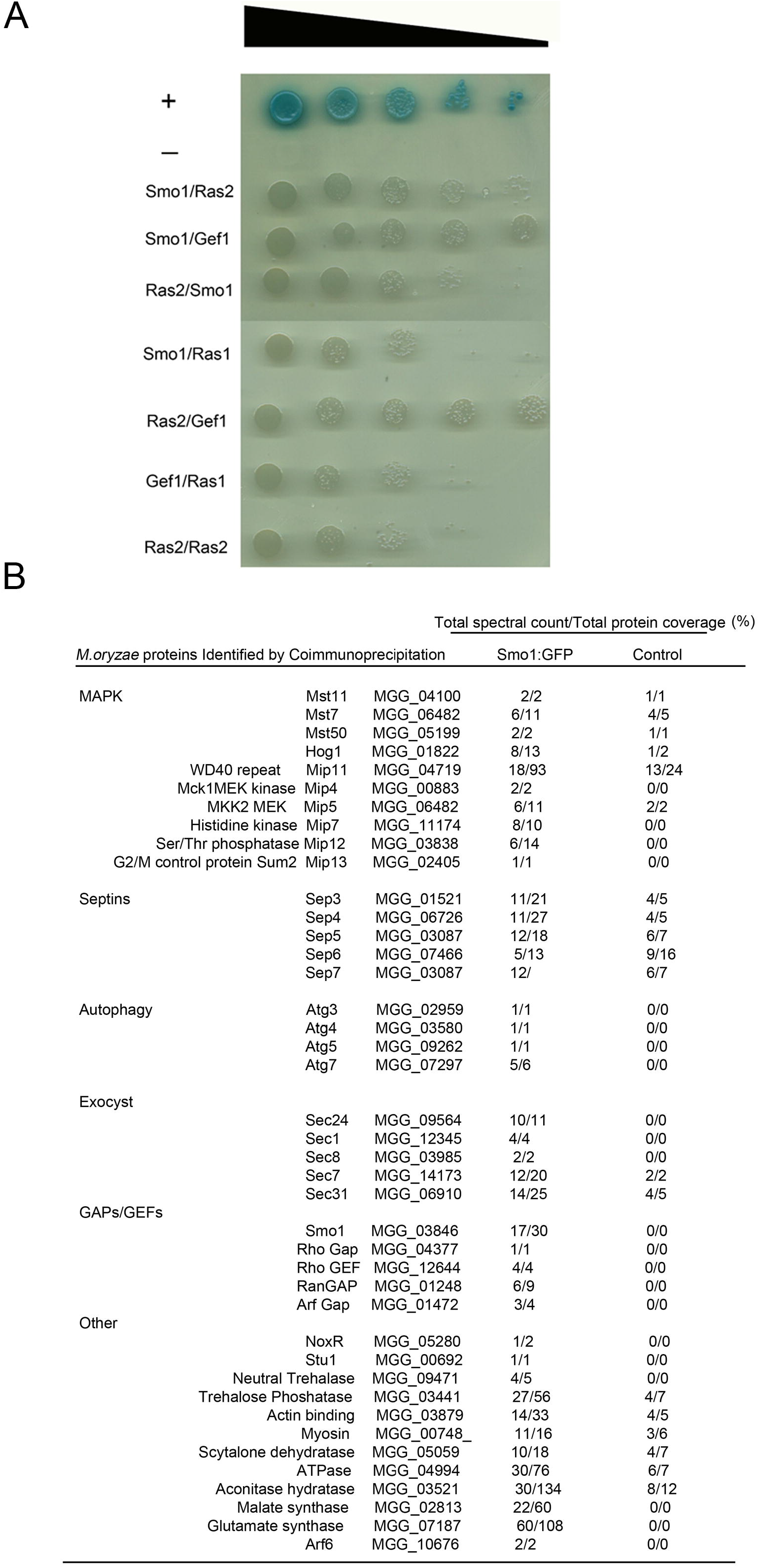
Yeast Two-hybrid analysis. **A**. Yeast two-hybrid screens were performed to determine putative physical interactions of Smo1 and to confirm its function as a Ras-GAP. *SMO1, RAS2,* and *RAS1* cDNAs were cloned into the bait vector pGBKT7, *SMO1, RAS2 and GEF1* cDNA were cloned into the prey vector PGADT7. Simultaneous co-transformation of the pGBK-Ras2 (bait–Ras2) and pGAD–Smo1 (prey–Smo1) vectors into the Y2H Gold strain results in the activation of all four reporter genes and growth on high stringency media (-His/-Ade/-Leu/-Trp/+X-α-Gal). Smo1/Gef1 showed the highest stringency interaction. (+ = positive control, – = negative empty prey vector control) **B**. Putative Smo1-Interacting Proteins identified in mycelial extracts of *M. oryzae* Smo1-GFP strain by co-immunoprecipitation with anti-GFP antibodies and mass spectrometry. Table shows spectral counts and protein coverage for each putative interaction.

Secondly, we carried out co-immunoprecipitation of Smo1-GFP from hyphae of *M. oryzae* and identified interacting proteins by mass spectrometry. This revealed interactions with the MAP kinase signalling pathway components, previously implicated in appressorium development, such as Mst11, Mst7 and Mst50, which all operate upstream of the Pmk1 MAP kinase, as well as the WD40 repeat protein Mip11, as shown in Figure 7B. Moreover, Smo1 interacts with the four core septins and with components of the exocyst complex, which are known to be associated with appressorium pore function (Dagdas *et al.* 2012; Gupta *et al.* 2015), as well as autophagy components that are also necessary for appressorium function (Kershaw and Talbot, 2009).

## Discussion

In this report, we have provided evidence that *SMO1* encodes a Ras GTPase-activating protein that plays important functions in cell shape determination and infection-related development in the rice blast fungus. *SMO1* is critical for rice blast disease and plays a significant role in conidium and appressorium shape determination and attachment to the leaf surface, in addition to an important regulatory function in the re-polarisation apparatus that operates within the appressorium. Smo1 is essential for septin recruitment and organisation at the appressorium pore, which in turn is necessary for F-actin re-organisation and penetration peg development (Dagdas *et al.* 2012). Smo1 physically interacts with Ras2, suggesting a model in which Ras signalling is required for appressorium repolarisation, as shown in Figure 8.

**Figure 8.**
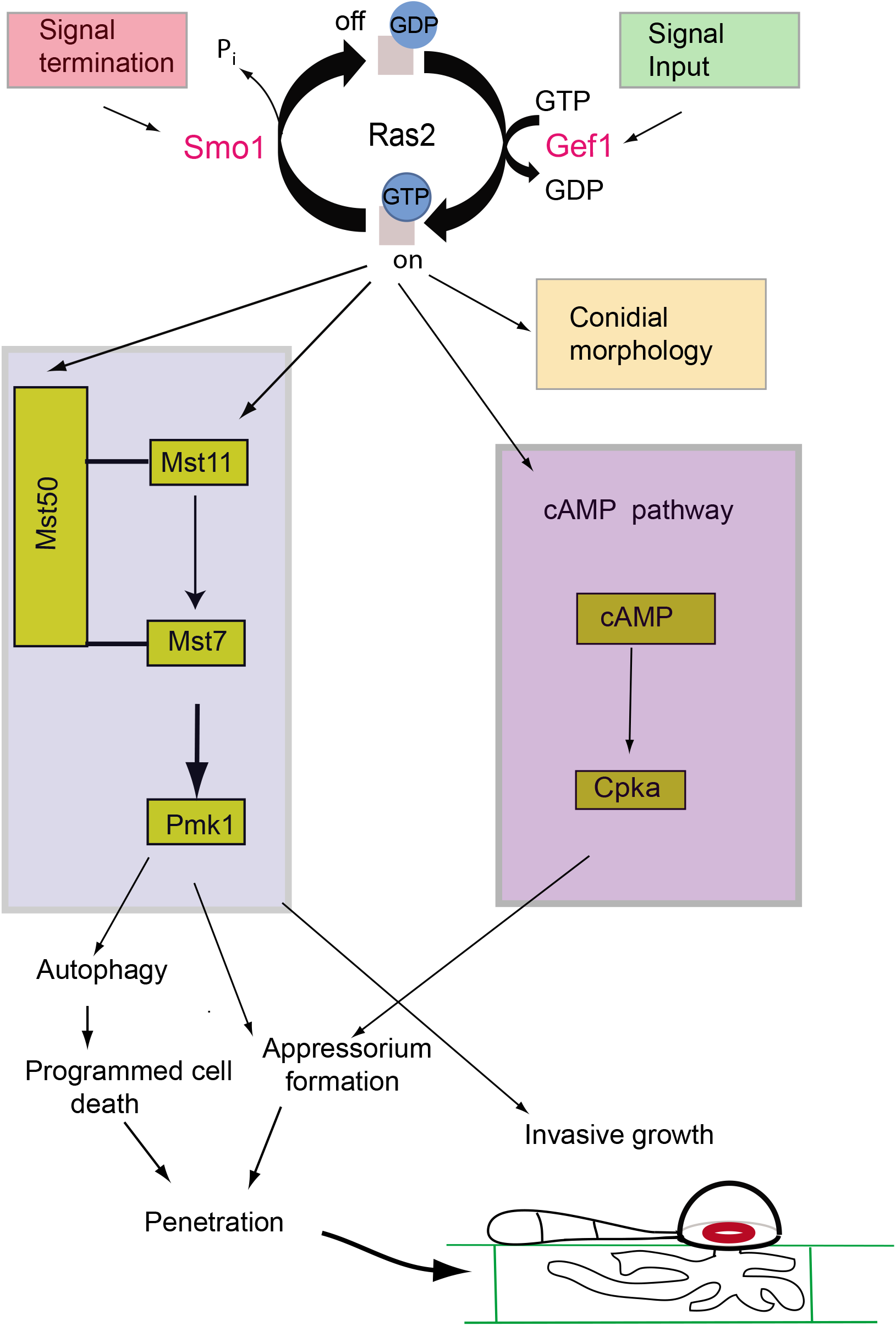
Model for the action of Smo1 in Ras2 signalling and its regulation of septin-dependent appressorium re-polarisation and plant infection.

Smo1^−^ mutants were so frequently identified during the early days of rice blast molecular genetic analysis that Hamer and colleagues (1989b) suggested that the *SMO* locus might be highly mutable. For example, isolation of spontaneous mutants with melanin pigment defects and benomyl-resistance identified double mutants that were also Smo1^−^ (CP665 and CP892). Separate genetic screens to identify mutants with a defect in appressorium development or a search for STM mutants with reduced capacity to attach to hydrophobic surfaces also mainly identified Smomutants (Hamer *et al.* 1989b). However, in spite of the rapid genetic mapping of the *SMO1* locus (Hamer and Givan, 1990), the gene proved to be extremely difficult to clone and after around 6 years of effort, *SMO1* cloning was finally abandoned. Recent advances in genome sequencing, the availability of numerous independent Smo1^−^ mutants, and the ability to carry out genetic crosses readily in *M. oryzae,* have now enabled us to identify *SMO1* and to understand the genetic events leading to frequent loss of gene function. Surprisingly, mutational events leading to inactivation of *SMO1* were all SNPs or small insertions/deletions in the coding sequence. This result contrasts with frequent deletion of the highly mutable *BUF1* melanin biosynthesis gene, which presumably occurs by transposon-mediated recombination (Chumley and Valent, 1991; Farman, 2002). The *SMO1* gene does not reside in a particularly transposon-rich region of the genome. Therefore, mechanisms for frequent isolation of Smo mutants and the original difficulties in cloning *SMO1* remain to be explained.

*SMO1* encodes a GTPase activating protein (GapA) involved in regulation of Ras proteins, and is one of four found in the *M. oryzae* genome. Ras-GA proteins work in conjunction with Ras-GEFs to regulate the activity of Ras proteins in response to external stimuli, affecting downstream signalling pathways necessary for regulation of morphological transitions necessary for growth in Eukaryotes (Boguski and McCormick, 1993). We have shown by yeast 2 hybrid analysis that Smo1 interacts with both Ras2 and Gef1, providing further evidence that Smo1 functions as a Ras-GAP and that it can form a complex with the corresponding guanine nucleotide exchange factor. Gap proteins are classified based on sequence homology within their Gap domains, with each domain specific for a class of G-proteins (Donovan *et al*. 2002). The different arrangements of domains, and in many cases the inclusion of distinct additional domains in the RasGAP family, suggest these proteins are subject to a diverse range of cellular interactions (Donovan *et al*. 2002). In *M. oryzae,* four putative RasGAPs can be classified into four different clusters according to their sequences and domain structures. Smo1 possess two domains, a GTPase activation domain for Ras-like GTPase and a RAS GAP C-terminal motif. These domains are characteristic of proteins belonging to the Ras-specific GAPs. The sequence placed Smo1 in a cluster with the *Aspergillus nidulans GapA* (Harispe *et al.* 2008) and *Gap1* from *Schizosaccharomyces pombe* (Imai *et al.* 1991). In *A. nidulans GapA* mutants exhibit abnormal conidiophores, delayed polarity maintenance characterised by apical swelling, and sub-apical hyphal branching (Harispe *et al.* 2008). In addition, F-actin distribution is lost in *ΔgapA* cells suggesting a role for *GapA* in F-actin cytoskeleton organisation, required for hyphal growth (Harispe *et al.* 2008). Mutation of Gap1 in S. *pombe* results in hypersensitivity to mating factor pheromone and the inability to perform efficient mating, which are identical phenotypes to those caused by activated ras1 mutations (Imai *et al.* 1991). The defect in polar growth suggests that *GAP1* is involved in polarity maintenance, working antagonistically with Ste6 in the regulation of Ras-GTPase in *S. pombe*. Involvement of a Ras-GAP in fungal morphogenesis was first reported for the basidiomycete white rot fungus, *Schizophyllum commune* in which *Gap1* deletion was shown to affect sexual development with mutants unable to form gills on fruiting bodies and producing no basidiospores. In addition, growth phenotypes suggested involvement in the maintenance of polarity (Schubert *et al.* 2006). The Smo1 mutant phenotypes observed are therefore consistent with those of GAP genes identified in other fungi, with effects on cell shape determination, polar/non-polar growth transitions, and regulation of the F-actin cytoskeleton.

Three other putative RasGAPs predicted in *M. oryzae,* MGG_11425.6 and MGG_08105.6 and MGG_03700.6, have not yet been characterised. MGG_03700 is a homolog of the *Saccharomyces cerevisiae* Iqg1, an essential gene shown by depletion and over-expression analysis to be required for cytokinesis and actin-ring formation (Epp and Chant, 1997). Iqg1 possesses a calponin-homology (CH) domain and IQ repeats in addition to the RAS GAP C-terminal motif (Epp and Chant, 1997). MGG_08105 is a homolog of the S. *cerevisiae* GAP *BUD2* which stimulates hydrolysis of the Ras2 homolog *BUD1.* Mutants defective in *BUD2* display random budding but no obvious growth defect (Park *et al.* 1993). MGG_11425 is a homolog of the S. *cerevisiae* RasGAPs *IRA1* and *IRA2,* which are negative regulators of Ras-cAMP signaling pathway required for reducing cAMP levels under nutrient limiting conditions (Tanaka *et al.* 1989; Tanaka *et al.* 1990).

Ras proteins are low molecular weight monomeric G-proteins which localise to the plasma membrane (Wennerberg *et al.* 2005) and switch between the active GTP-bound and inactive GDP-bound status, competitively regulated by GEF’s and GAPs (Boguski and McCormick, 1993). Ras proteins have intrinsic GTPase and GDP/GTP exchange activity, but GAP and GEF proteins work to effect a more tightly regulated process. In S. *cerevisiae,* the two Ras proteins, Ras1 and Ras2 are both essential for growth and both function to activate adenylate cyclase (Tamanoi, 2011). The RAS/cAMP/PKA pathway in S. *cerevisae* regulates a variety of processes, including cell cycle progression and life span (Tamanoi, 2011). *M. oryzae* also has two Ras-encoding genes, *MoRAS1* and *MoRAS2,* and both have been characterised. In the *Δras1* deletion mutant no distinct phenotypes were observed other than a slight reduction in conidiation (Zhou *et al.* 2014). *RAS2,* however, is thought to be an essential gene and has only been characterised by generation and expression of a *Ras2* dominant active allele. *MoRAS2^G18V^* transformants formed morphologically abnormal appressoria on both hydrophilic and hydrophobic surfaces, suggesting that dominant active *RAS2* can bypass surface attachment requirements for appressorium formation (Zhou *et al.* 2014). *MoRAS2^G18V^* showed increased Pmk1 phosphorylation and elevated cAMP levels in aerial hyphae. cAMP-PKA signalling has been shown to be important for initial surface recognition and appressorium generation and for generation of turgor pressure necessary for infection. The cAMP-dependent PKA mutant, *ΔcpkA,* produces long germ tubes and small non-functional appressoria (Xu *et al.* 1997), whilst the mitogen–activated protein (MAP) kinase Pmk1 is essential for appressorium formation and invasive growth (Xu and Hamer, 1996) In both *Δpmk1* and *Δcpka* mutants, expression of *MoRAS2^G18V^* had no effect on appressorium morphogenesis, suggesting that Ras2 functions upstream of both cAMP and Pmk1 signalling pathways (Zhou *et al.* 2014). Deletion of several upstream components of the Pmk1 pathway, including *MST50, MST11* and *MST7* result in defects in appressorium development and plant infectction (Park *et al.* 2006; Zhao *et al.* 2005; Zhou *et al.* 2014). Mst50 functions as an adapter protein of the Mst7-Mst11-Pmk1 cascade involved in activating Pmk1 in *M. oryzae* (Zhao *et al.* 2005). Both Mst50 and Mst11 have been shown to interact with MoRas1 and MoRas2, by yeast two-hybrid assays (Park *et al.* 2006) and deletion of the Ras-association (RA) domain of Mst11 blocked Pmk1 activation and appressorium formation (Qi *et al.* 2015), supporting a role for Ras signalling in activation of the Pmk1 pathway. It has been shown recently that the transmembrane mucins, Msb2 and Cbp1, function together to recognise extracellular signals through Ras2 (Wang *et al.* 2015). Pmk1 phosphorylation was reduced in a *Momsb2* mutant but blocked in a *Momsb2 cbp1* double mutant, which was non-pathogenic.(Wang *et al.* 2015). Affinity purification was used to identify a series of Mst50-interacting proteins (MIPs) as well as upstream kinases of the Mps1 pathway, and also the histidine kinase Hik1 (Li *et al.* 2017). These interactions suggest a role for Mst50 in three different signalling pathways. However, domain deletion experiments showed that the Mst50 Ras-association domain was not important for response to oxidative stress (Li *et al.* 2017). *M.oryzae* Ras2 is therefore essential for cellular viability, and a key mediator between both the Pmk1 MAPK and cAMP signalling cascades (Qi *et al.* 2015; Zhou *et al.* 2014). Activation and deactivation of Ras2 regulates developmental switches/pathways necessary for growth and pathogenicity of the fungus (Zhou *et al.* 2014).

When considered together, our results suggest that Smo1 acts as a negative regulator of Ras2 and this is why *Δsmo1* mutants display such severe developmental defects, including mishappen spores and appressoria, long germ tubes and a failure in penetration peg development (see model in Figure 8). These phenotypes point to a defect in the maintenance of polarity that is required for morphological transitions in the fungus. These developmental effects are a consequence of the disruptions to both septin and F-actin dynamics in *Δsmo1* mutants, which are essential for plant infection. SMO1-dependent regulation is therefore required for the morphological transitions, and cell shape generation processes that are associated with asexual reproduction and plant infection by the blast fungus.

## Acknowledgements

This work was funded by a Biological Sciences and Biotechnology Research Council (BBSRC) Industrial Partnership Award with Syngenta (BB/) and a European Research Council Advanced Investigator Award to NJT under the European Union’s Seventh Framework Programme *(FP7/2007-2013)* / ERC grant agreement n° 294702 GENBLAST. This is contribution number 18-374-J from the Kansas Agricultural Experiment Station. This paper is dedicated to John Hamer and the brave students and postdocs in his research group who tried so hard to clone *SMO1* in the early 1990’s; including Kathy Dobinson, Scott Givan, Steve Harris, Michelle Momany, Yankyo Salch, and Verel Shull. The corresponding author (NJT) shares your pain, having spent much of 1991 also trying in vain. The authors all salute you.

---

**Table S1** Smo1 mutants used in this study and characteristics of the genome sequences generated

**Table S2** Primers used in this study

**Figure S1**. Multiple sequence alignment using Clustal X (Larkin et al. 2007) showing GTPase activating proteins from *M. oryzae* in Maximum likelihood phylogenetic tree created using PhyML. Bootstrap support values of 70 or greater are indicated on tree.

**Figure S2**. Complementation of the *Δsmo1* mutant restores wild type phenotypes. (A) Micrographs showing spore morphology of Guy11 and *Δsmo1* mutant complemented with the Smo1-GFP fusion construct, and the *smo1^−^* strain CP788 complemented with wild type *SMO1* allele. Bar = 10 μm (B) Photographs of leaves from infected seedlings of rice cultivar CO-39 inoculated with conidial suspensions (1 × 10^5^ ml^−1^) of wild type, *Δsmo1* and *Δsmo1* mutant complemented with *SMO1-GFP.*

**Figure S3**. (A) Cellular localisation of H1-RFP by live cell imaging in the *Δsmo1* mutant visualised by epifluorescence microscopy over a time course of infection related development. Bar = 10 μm. (B) Calcofluor-white staining of wild type and *Δsmo1* mutant at 24 hpi. Bar = 10 μm.

